# Dual specificity phosphate 1 (DUSP1) as a non-invasive circulating biomarker candidate in preeclampsia

**DOI:** 10.1101/2024.10.12.618015

**Authors:** Jonatane Andrieu, Agathe Donet, Jean-François Cocallemen, Guillaume Charbonnier, Noémie Resseguier, Julien Paganini, Jean-Louis Mège, Soraya Mezouar, Florence Bretelle

## Abstract

**Background:** Preeclampsia (PE) is a multisystem pregnancy complication constituting a major cause of maternal and fetal morbidity and mortality. Factors pointing to a placental origin are the development of the pathology only during pregnancy, and its disappearance in the post-partum period.

**Methods:** Here, we aim to identify new early predictive biomarkers based on a transcriptional signature of PE using RNAseq. Whole blood and serum samples were collected at the time of the first event of PE (V1) and same samples after remote delivery (30-60 postpartum days, V2). These two samples enabled investigation of PE markers found in V1 but absent in V2. To confirm that these candidates are associated with PE, an investigation of associated placental biopsy was also realized (J0).

**Results:** Our study identified a specific signature of PE including five Gene Ontology clusters including “angiogenesis and differentiation”, “cell cycle”, “cell adhesion”, “inflammatory response” and “cellular metabolism”. Interestingly, *DUSP1* gene was found specifically modulated in PE. Pregnant women with PE have a higher concentration of DUSP1 in serum compared to healthy donors. Interesting, at a distance from childbirth (V2), DUSP1 finds a rate like the control group showing the predictive interest of DUSP1 as a promising predictive biomarker of PE.

**Conclusions:** The investigation of DUSP1 in a prospective study with a larger cohort, including the severity aspect of the disease, is necessary to confirm its value as a predictive biomarker in PE.

## Introduction

Preeclampsia (PE) is a progressive, multisystem pregnancy complication that affects 3 to 5% of pregnancies, making it one of the major causes of maternal and fetal morbidity and mortality 1. PE is responsible for hematological complications, as well as organs failure that can be severe, notably placental, neurological, hepatic, pulmonary, renal and cardiovascular 2,3. Fetal complications include life-threatening complications such as intrauterine growth retardation, malformation and induced prematurity 4. PE is a complex pathological process that originates at the mother-fetal interface 5,6. It is accepted that PE is a disease of the maternal endothelium with placental origins. Factors pointing to a placental origin are the development of the pathology only during pregnancy, and its disappearance in the post-partum period.

The use of an early predictive marker of PE is crucial. A number of markers were initially identified, notably the combination of several early markers - clinical (mean arterial pressure), ultrasound (uterine artery pulsatility index) and biological (pregnancy associated plasma protein (PAPP-A) and placental growth factor (PlGF)) - to predict the risk of preterm PE, with a positive predictive value of around 10 to 15% meaning 90%, for the fetal medicine foundation (FMF) test, that it will get a high-risk result but not end up developing preterm PE^7^. Other studies have focused on the trophoblastic cells, as placental cells, looking at their processes of migration and invasion ^8^. Markers such as programmed death-ligand 1 (PD-L1) ^9^ and angiopoïetine like 4 (ANGPTL4) ^10^ have been shown to significantly increase trophoblast invasion and migration in PE, as well as implicating the yes-associated protein (YAP)-Hippo trophoblast differentiation pathway ^11^. However, these factors only provide a better understanding of the physiopathology of the PE.

There has been growing interest in early predictive biomarkers for PE. Effective predictive tests will facilitate early diagnosis, targeted monitoring and prompt management, using biomarker(s) capable of predicting early pregnancy (less than 16 weeks) in women at high risk of clinical complications in the prevention of PE ^12^. The anti-angiogenic factor, soluble fms-like tyrosine kinase 1 (sFlt-1), found in the placenta and measured in plasma and serum, was proposed as a specific biomarker to the onset and severity of PE ^13^. The evaluation of the ratio of sFlt-1 and the pro-angiogenic factor, PlGF, were found to present a negative predictive value ^14^ and can be used to predict the short-term absence of PE in women in which the disease was previously suspected clinically. Unfortunately, the predictive value is strongly linked with prevalence of the disease. Ongoing studies are focused on the selection of women to prevent PE onset with acid salylic prescription. ASPRE trial showed that identification of at risk women using a score including mean arterial pressure, uterine-artery pulsatility index, and maternal serum PAPP-A and PlGF reduce early PE ^15,16^. The overall rate was not decreased which encourage further studies on the identification of new tools or factors.

The aim of this study was to identify new early biomarkers of PE based on a transcriptional signature of PE at the time of the event, using both maternal blood (peripheral) and placental biopsy (local). The secondary objective was to evaluate the evolution of the expression of this signature during pregnancy progression, particularly at the time of delivery (samples from maternal blood (peripheral) and placental biopsy (local)). Finally, this study aims to investigate the disappearance of expression of these markers at distance from the event in the postpartum period.

## Material and Methods

### Ethics statement

This single-center prospective cross-sectional study was conducted in accordance with the Declaration of Helsinki and the French law on research involving humans. The protocol of the study was approved by an independent national review board ethics committee “CPP Sud Mediterranean 1” n° 2010-A00633-36. All pregnant women provided written informed consent and were recruited at the gynecology-obstetrics department of the “Hôpital de la Conception” and “Hôpital Nord” (Marseille, France) from February 2019 and July 2020.

### Study participants and sample collection

Pregnant women consisting of 10 healthy controls and 10 patients who suffered of PE at 20 and 37 gestational ages were included (**Table 1**). PE pregnant women presented arterial hypertension (systolic blood pressure greater than or equal to 140 mmHg and/or diastolic blood pressure greater than or equal to 90 mmHg) associated with positive proteinuria (positive urine dipstick or proteinuria greater than 0.3 g per 24 hours) during pregnancy. PE and control groups were matched for patient age and gestational age at inclusion. Clinical parameters were recorded including maternal age, geographical origin, body mass index (kg/m^2^) as well as obstetrical characteristics (gestational age, parity, induced or spontaneous pregnancy, associated gravidic pathologies). Detailed fetal outcome was also monitoring such as ultrasound findings, fetal heart rate analysis and neonatal data. Total blood sample (PAXgene tube, PreAnalytiX) was taken at the time of PE diagnosis (and matched on gestational age at inclusion for controls) and was repeated 4 to 6 weeks postpartum (**Figure S1**). PAXgene tubes were stored at 4°C for 24 hours, before freezing for 24 hours at -20°C and then permanently stored at -80°C. At-term placental biopsy was also performed at the time of delivery for each of the included women. Placental biopsy was defined as a macroscopically placental area of 2x2 cm including both chorionic and basal membranes. Biopsies were immediately preserved in RNA*later* (Thermo Fisher Scientific) for 24 hours at 4°C, before freezing for 24 hours at -20°C, then permanently at -80°C.

**Table 1.** Initial characteristics of the population at the time of inclusion. BMI: body mass index. HELLP syndrome: syndrome of hemolysis, elevated liver enzymes, and low platelet

|  | Control (n=10)<br>n (%) | Preeclampsia<br>(n=10)<br>n (%) | *p-value* |
| --- | --- | --- | --- |
| *Pregnant women* |  |  |  |
| Age (years) | 29.50 ± 4.45 | 31.20 ± 7.43 | 0.54 |
| Geographical origin |  |  | 0.58 |
| • Caucasian | 7 (70) | 9 (90) |  |
| • African | 3 (30) | 1 (10) |  |
| • Asian | 0 | 0 |  |
| BMI (Kg/m<sup>2</sup>) | 23.70 ± 4.27 | 24.50 ± 3.98 | 0.68 |
| Smoking status |  |  | - |
| • Absence | 9 (90) | 10 (100) |  |
| • Active (>10 cig/day) | 1 (10) | 0 (0) |  |
| *Obstetrical characteristics* |  |  |  |
| Gestational age at diagnosis (weeks) | 31.4 ± 4.30 | 29.5 ± 3.13 | 0.29 |
| Conception type |  |  | 0.47 |
| • Spontaneous pregnancy | 10 (100) | 8 (80) |  |
| • Induced IVF pregnancy | 0 (0) | 2 (20) |  |
| Delivery route |  |  | 0.0007 |
| • Vaginal delivery | 8 (80) | 0 (0) |  |
| • Caesarean section | 2 (20) | 10 (100) |  |
| Maternal complications |  |  | 0.0867 |
| • Absence | 10 (100) | 6 (60) |  |
| • Presence | 0 (0) | 4 (40) |  |
| · Uncontrolled hypertension | 0 (0) | 3 (30) |  |
| · Proteinuria >6g/day | 0 (0) | 2 (20) |  |
| · Acute renal failure | 0 (0) | 1 (10) |  |
| · HELLP syndrome | 0 (0) | 2 (20) |  |
| *Fetal outcome* |  |  |  |
| Gestational age at birth | 39.5 ± 1.13 | 30.1 ± 3.1 | >0.001 |
| Time between inclusion and delivery (days) | 61.6 ± 31.25 | 3.3 ± 3.05 | >0.001 |
| Fetal growth |  |  | 0.21 |
| • Eutrophic fetus | 10 (100) | 7 (70) |  |
| • Intrauterine growth retardation | 0 (0) | 3 (30) |  |
| Fetal Doppler |  |  | 0.21 |
| • Normal fetal Doppler | 10 (100) | 7 (70) |  |
| • Doppler anomalies | 0 (0) | 3 (30) |  |
| Neonatal complications |  |  | 0.02 |
| • Absence | 8 (80) | 2 (20) |  |
| • Presence | 2 (20) | 8 (80) |  |
| · Fetal growth restriction | 0 (0) | 4 (40) |  |
| · Respiratory distress | 2 (0) | 3 (30) |  |
| · Neonatal death | 0 (0) | 1 (10) |  |
| Birth weight | 3174.4 ± 281 | 1203.5 ± 611.1 | >0.001 |

### RNA extraction

RNAs from total blood samples were extracted using the PAXgene Blood RNA kit according to the procedure recommended by the manufacturer (Qiagen). Briefly, total blood was lyzed using proteinase K and the nucleic acids were precipitated by the addition of ethanol. DNA was digested with DNase I RNase-free for 15 minutes at room temperature. Total RNAs were eluted and then incubated at 65°C for 5 minutes before being placed at -80°C.

Total RNAs from placenta biopsies were extracted using the RNeasy Mini Kit according to the procedure recommended by the manufacturer (Qiagen). After dissolution of placental tissue with RLT-β-mercapto-ethanol, nucleic acids were precipitated with the addition of ethanol. DNA was digested with DNase I RNase-free for 15 minutes at room temperature. Total RNAs were eluted before being placed at -80°C.

The quality and quantity of the extracted RNAs were evaluated using the Bioanalyzer 2100 (Agilent Technologies) and by NanoDrop Spectrophotometer (Nanodrop Technologies).

### RNA-sequencing and data processing

Reads were aligned and counted with STAR (https://doi.org/10.1093/bioinformatics/bts635) on the hg19 assembly with the GENCODE v19 annotations. The raw gene count table was variance-stabilized and reduced into principal components and UMAP for quality control. The raw count table was also used to compute differential expression analysis (DEA) using Deseq2 framework ^17^. Finally individual DEA results were gathered into integration plots. Data from RNASeq data analysis were submit on the GEO data collection (GSE262147).

### Quantitative reverse transcription-polymerase chain reaction (qRT-PCR)

Reverse-transcription of isolated RNA was performed using a Moloney murine leukemia virus-reverse transcriptase kit (Life Technologies) and oligo (dT) primers. The expression of genes was evaluated using real time qPCR, Smart SYBR Green fast Master kit (Roche Diagnostics) and specific primers (**Table S1**). The qPCRs were performed using a CFX Touch Real-Time PCR detection system (Bio-Rad). The results were normalized by the expression of the *ACTB* housekeeping gene and are expressed as relative quantity (RQ) of investigated genes with 2^-ΔCt^ with ΔCt = Ct_Target_ – Ct*_ACTB_* as previously described ^18^.

### Immunoassays

FLT1 and DUSP1 levels were quantified in serum from study population with appropriate immunoassays according to the manufacturer’s instructions (Antibodies). The sensitivity was 6.99 pg/ml for FLT1 and 9.4 pg/ml for DUSP1.

### Protein interactome

The protein interactome between DUSP1 and FLT1 was created using String functional association networks protein software.

### Statistical analysis

The statistics of the initial characteristics of the population and from the fat were carried out with the software R 3.6.1. The variables were described by the mean and standard deviation. The qualitative variables were described by their percentage and p-value. Categorical variables were compared by a chi2 test or by a Fisher exact test. The alpha risk was defined at 5%.

Statistical analysis of gene signatures was performed with GraphPad Prism 6 (Graphpad Software Inc.). Gene expression was analyzed using the one-way ANOVA test and Tukey’s multiple comparisons test. Values represent the mean ± standard deviation. The limit of significance was set up at *p<0.05*.

### Results Study design

We conducted a prospective cross-sectional study to investigate novel biomarkers for PE diagnosis. Ten patients with PE were included during the study period in a university medical center. Ten patients with normal pregnancy and devoid of significant antecedents were matched as controls to previous patients, on maternal age and gestational age to the diagnosis of PE.

The scheme of the study was shown in **figure S1**. Whole blood and serum samples were collected at the time of the first event of PE (V1) and same type of samples after remote delivery (30 to 60 postpartum days, V2). These two samples enabled investigation of PE markers found in V1 but absent in V2. To confirm that these biomarkers are associated with PE, an investigation of associated placental biopsy collected after delivery was also realized (J0).

We first focused on the study population at the time of inclusion. As illustrated in the **table 1**, the maternal age at diagnosis was comparable in cases and controls, respectively 31 ± 7.43 years and 29.5 ± 4.45 years (*p=0.54*). Gestational age at diagnosis showed no significant difference between the two groups: 29.54 ± 3.13 gestational age in PE and 31.38 ± 4.30 gestational age in controls (*p=0.29*). No significant differences were also observed for the body mass index, smoking and for the severity of pregnancy.

Considering the outcomes of pregnancy of the two groups (**Table 1**), as expected the gestational age at delivery was significantly earlier in the PE group (30.07 ± 3.12) than in the control group (39.52 ± 1.13) (*p>0.001*). The time between inclusion and delivery was significantly shorter in the PE group (3.3 ± 3.05) than in the control group (61.6 ± 31.25) (*p>0.001*). Patients with PE presented birth by cesarean section in 100% of cases, compared to 20% in the control group (*p=0.0007*). Serious maternal complications were observed in the PE group, such as uncontrolled hypertension (30%), heavy proteinuria (20%), acute renal failure (10%) and HELLP syndrome (20%). However, no significant differences were observed between the two groups (*p=0.0867*). Similarly, there is a significant difference in neonatal outcomes. Neonatal weight was significantly lower in PE (1203.5 ± 611.1 *versus* 3174.4 ± 28, *p>0.001*). Neonatal complications are significantly increased in PE (80% *versus* 20%, *p=0.02*). In our cohort, they mainly consisted of severe sepsis (40%) and respiratory distress (30%), and neonatal death in 1 case.

### Preeclampsia RNA profile

After the raw data normalization, the differences between samples from PE pregnant women and healthy donors were represented in the **figure 1**. The clustering hierarchical heat map highlighted that placental samples were positioned in a separate branch of whole blood samples (**Figure 1A**). RNA-seq analysis revealed 23,919 differentially expressed genes (fold change > 2 and false-discovery rate (FDR) < 0.05) as illustrated in the volcano plot (**Figure 1B**). Principal component analysis demonstrated contrasts among the two investigated group regarding the type of sample (**Figure 1C**), but not considering the investigated groups (**Figure 1D**). When we excluded the sample variable, we did not distinguish grouping in terms of group (control and PE) for the different individuals investigated (**Figure 1E, 1F**).

**Figure 1.**
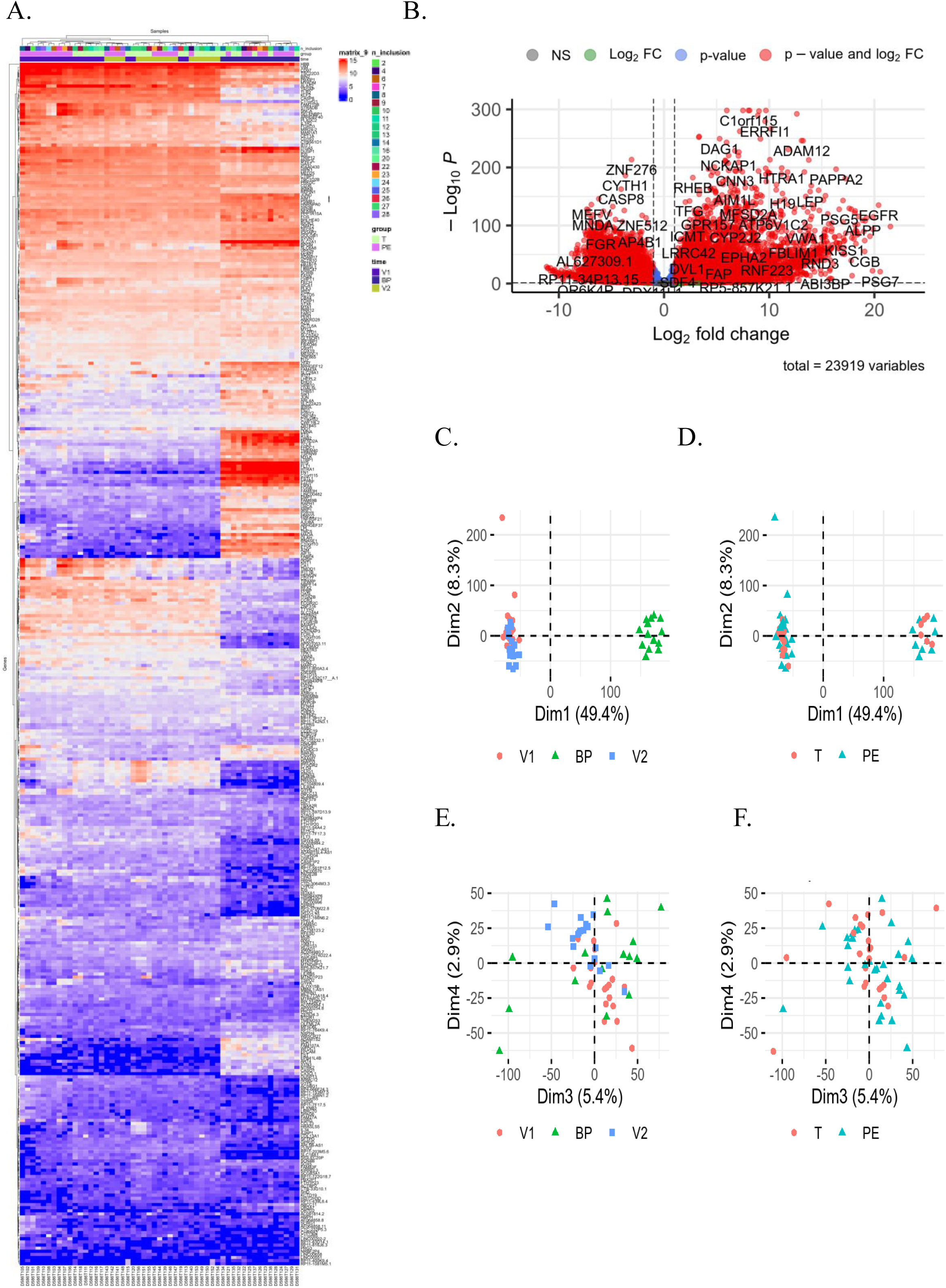
RNAseq data. (**A**) Hierarchical clustering and (**B**) volcano plot highlighted modulated genes from RNAseq data analysis that revealed 23,919 differentially expressed genes (fold change > 2 and false-discovery rate (FDR) < 0.05). (**C - F**) Principal component analysis illustrated the repartition of investigated groups (V1, BP and V2). BP, Biopsy from Placenta; FC, Fold Change.

We next investigated gene modulation in PE and control groups. After adjustment and selection with variance and *p<0.05* we consistently identified 300 genes modulated. Focusing on the time covariate we observed 27% genes up-modulated (81) and 20% down-modulated (61) genes **(Figure 2A**). Interestingly, at the time of the first inclusion (V1), corresponding to the first event of PE, 108 genes were found to be modulated upwards in whole blood from PE group compared to controls (**Figure 2B**). This high rate of up-modulated genes was also found in the transcriptional signature of the placental biopsy (**Figure 2C**). In contrast, for V2, corresponding to the patient sample taken at distance from childbirth, the number of up- and down-modulated genes was similar. The aim of this study was to determine relevant biomarkers that might reflect the pathophysiology driving PE. Thus, we interested in genes modulated in whole blood in V1, absent in V2 and present in the placenta for the PE group compared to the control group. In this condition 24 genes, as a specific signature of PE, were identified as illustrated in the hierarchical clustering (**Figure 3A**) and volcano plot (**Figure 3B**). Gene Ontology (GO) analysis considering “Biological Process” revealed five GO clusters as follow. In ascending order, we identify 30.8% of genes associated with “angiogenesis and differentiation”, 26.9% with “cell cycle”, 19.2% with “cell adhesion”, 15.4% with “inflammatory response” and 7.7% with “cellular metabolism” (**Figure 3C** and **Table S2**).

**Figure 2.**
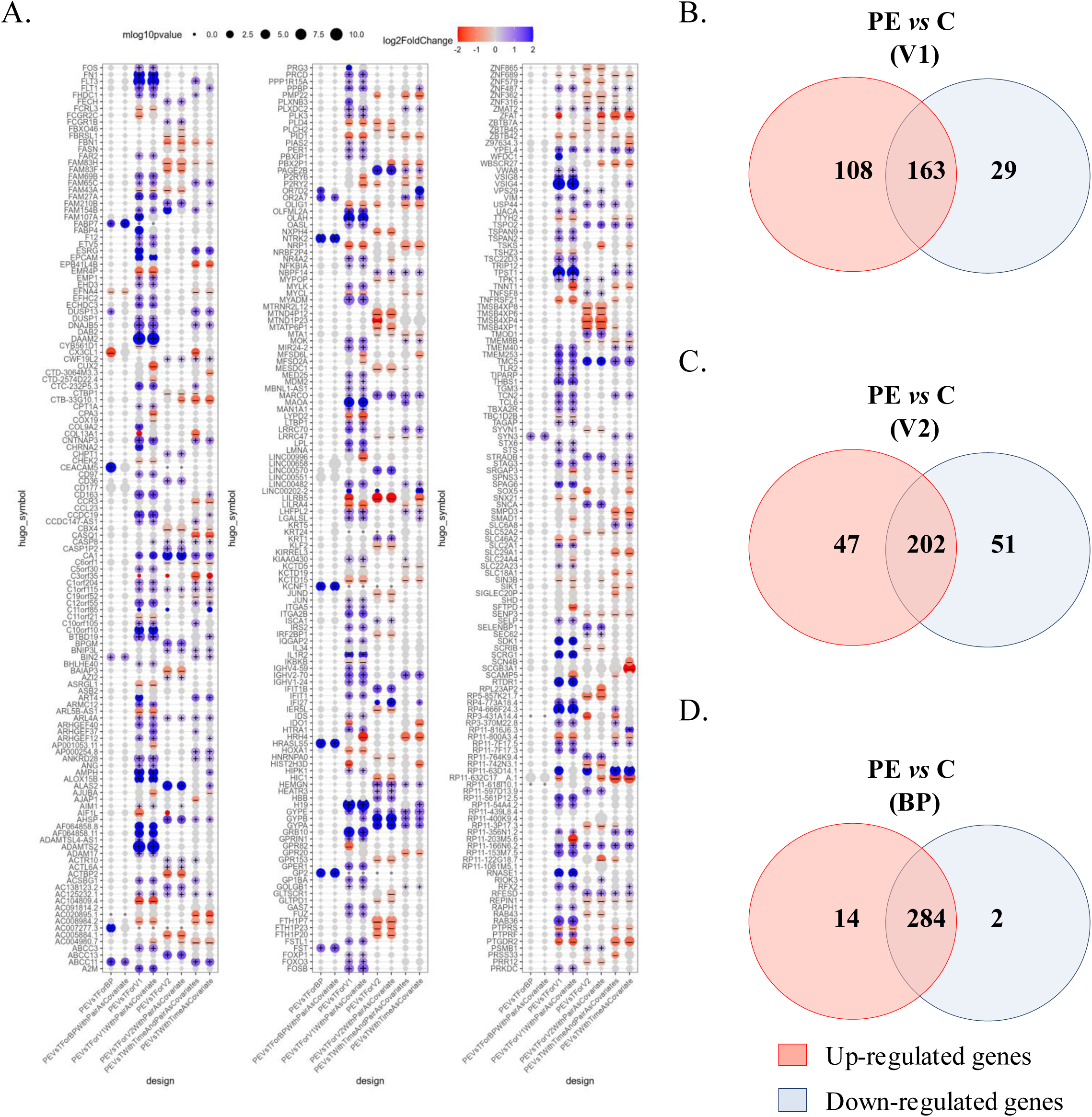
**Modulated genes associated to preeclampsia** (**A**) Clusters of 300 modulated genes obtained after adjustment and selection with variance and *p<0.05* value. (**B**) Venn diagrams illustrated up- and down-modulated genes for preeclampsia (PE) *versus* control (C) group for V1, V2 and BP. BP, Biopsy from Placenta.

**Figure 3.**
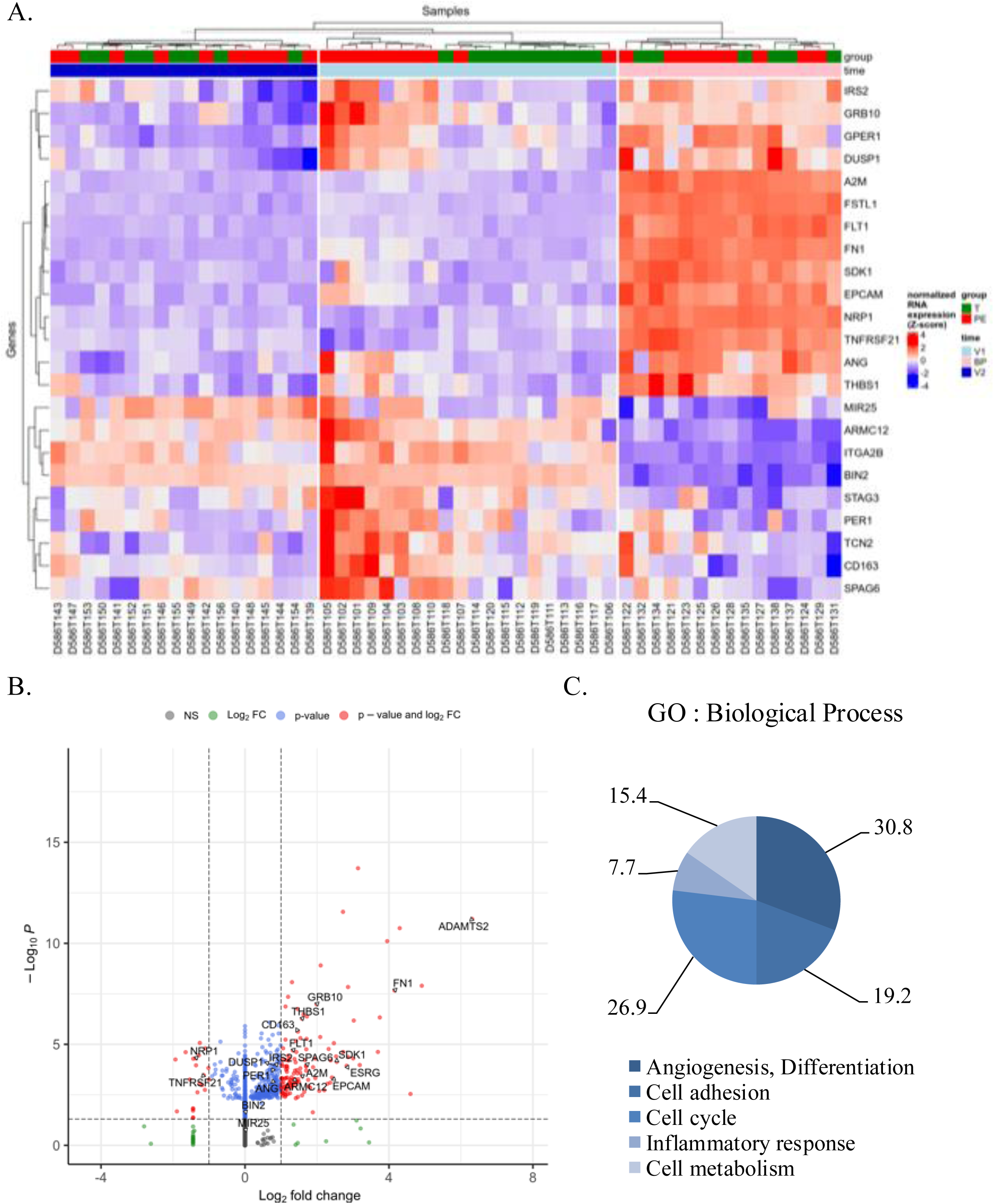
**Specific genes associated to preeclampsia**According to the selection of genes of interest modulated in V1, absent in V2 and present in the biopsy from placenta (BP) for the PE group compared to the control (T) group we obtained 24 genes as a specific signature of PE. (**A**) Hierarchical clustering and (**B**) volcano plot illustrated the 24 modulated genes. (**C**) Graph illustrating the Gene Ontology (GO) analysis considering “Biological Process” including the percentage of genes associated with “angiogenesis and differentiation”, “cell cycle”, “cell adhesion”, “inflammatory response” and “cellular metabolism”. FC, Fold Change.

### Identification of a specific signature for preeclampsia

Genes identified were next evaluated using quantitative-retro-transcriptase polymerase chain reaction (**Figure 4**). Among genes associated to the “cellular metabolism” only *A2M* presented a significant increased on time V1 in PE compared to control group (*p=0.0236*) but without differences between V1 and V2 for PE group (**Figure 4A**). No significant differences were observed for *TCN2, SPAG6* and *ADAMTS2* genes. Also, in the “inflammatory response” only *TNFRSF21* was found significant increased on time V1 in PE compared to control group (*p<0.0001*) and for PE group a significant decreased was observed between V1 and V2 (*p<0.0001*) (**Figure 4B**). In this cluster, no differences were observed for *CD163*. Among the 4 genes associated with the cluster “cell adhesion” (*ITG12B*, *THBS1*, *EPCAM*, *SDK1*), two genes (*THBS1*, *SDK1*) were found differently modulated according to PE and control groups (**Figure 4C**). *THBS1* was increased in PE group at the placenta level (*p=0.0073*) without statistically significance at the blood level. *SDK1* was found significant increased on time V1 in PE compared to control group (*p=0.0099*) and for PE group a significant decreased was observed between V1 and V2 (*p=0.0267*) (**Figure 4B**). Among the 7 genes associated to the cluster “cell cycle” (*BIN2*, *PER1*, *MIR25*, *IRS2*, *ESRG*, *STAG3*), 3 genes were found differently modulated according to PE and control groups (**Figure 4D**). At the blood level, *MIR25* found significant increased on time V1 in PE compared to control group (*p=0.0012*) and for PE group a significant decreased was observed between V1 and V2 (*p=0044*) (**Figure 4D**). At the placental level, *ESRG* and *GPER1* presented a significant increase in PE compared to control group (*p<0.0001* and *p=0.0017*, respectively). Finally, we found 8 modulated genes associated to the “angiogenesis and differentiation” cluster (*GRB10, FN1, FLT1, DUSP1, NRP1, ANG, ARMC12, FSTL1*) (**Figure 4E**). Among them, 6 genes were found differentially modulated among the investigated groups (*FN1, ANG, GRB10, FSTL1, DUSP1*). Three genes were modulated either at the blood or placenta level. *FN1* and *ANG* were significantly increased in PE compared to control from placenta biopsies (*p=0.0007* and *p<0.0001*, respectively). *FSLT1* and *GRB10* increased on time V1 in PE compared to control group (*p=0.0037* and *p<0.0001*, respectively) and for PE group a significant decreased was observed between V1 and V2 (*p<0.0001* and *p=0007*, respectively). Interestingly, in our study we found the *FLT1* gene, whose interest as a biomarker in PE was well-documented ^14^. In our study scheme, we observed that *FLT1* is significantly increased in V1 in the PE group compared to the control group (*p<0.0001*). Also, focusing on the PE group, *FLT1* decreased in V2 compared to V1 (*p<0.0001*). Locally, in placental biopsy of women with PE *FLT1* was significantly high expressed compared to controls (*p<0.0001*) confirming that *FLT1* was a biomarker of interest in PE pathophysiology as previously described ^14^. Compared to all the investigated genes, *DUSP1* showed the same state of significant expression modulation as *FLT1*: (1) increased in V1-PE compared to V1-Control (*p=0.0185*), (2) decreased in V2-PE compared to V1-PE (*p=0.0011*) and (3) increased in placental level in PE compared to control group (*p=0.0006*). Taking together, our study highlights the interest for the *DUSP1* as a specific gene of PE.

**Figure 4.**
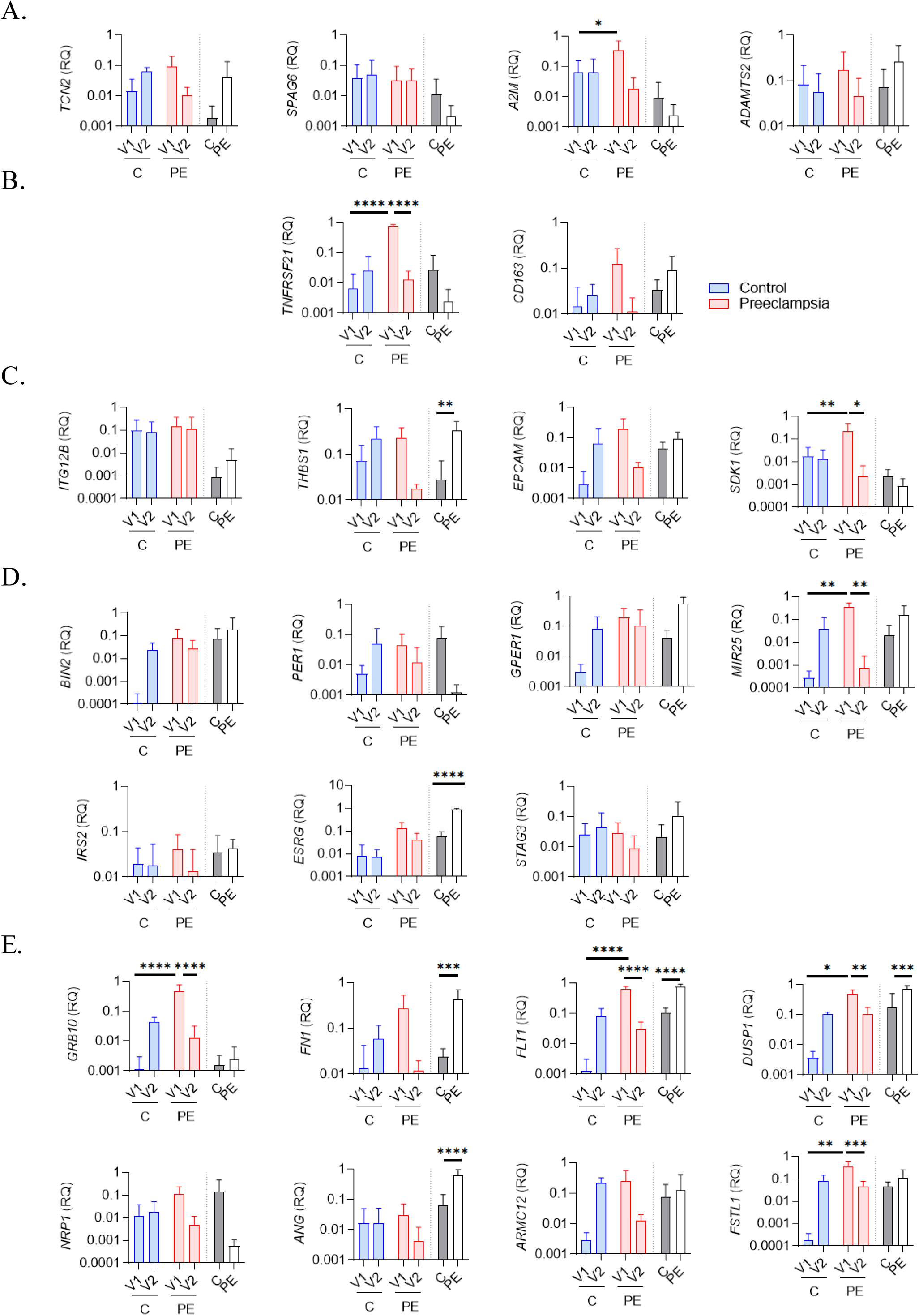
qRTPCR evaluation of specific genes associated to preeclampsia. Relative quantity evaluation of genes involved in (**A**) “cellular metabolism”, (**B**) “inflammatory response”, (**C**) “cell adhesion”, (**D**) “cell cycle” and (**E**) “angiogenesis and differentiation” pathways. Modulated genes were obtained after qRTPCR experiments using whole blood (V1, V2) and biopsy from placenta (BP) from 6 healthy (C) and 6 preeclamptic women (PE). Data values represent the mean ± SEM whose experiments were carried out in triplicate. Statistical analysis was performed with one-way ANOVA and Tukey’s multiple comparison test. \**p*≤*0.05*, \*\**p*≤*0.01*, \*\*\**p*≤*0.001* and \*\*\*\**p*≤ *0.0001*.

### *DUSP1* modulation in preeclampsia

We next evaluated levels of DUSP1 in plasma samples using immunoassays. As illustrated in the **figure 5A**, DUSP1 was barely detected in healthy donor plasma during pregnancy (V1) or post-partum (V2). Interestingly, pregnant women with PE have a higher concentration of DUSP1 in V1 compared to controls (*p<0.0001*), suggesting that DUSP1 could be an interesting biomarker. Focusing on the PE group we showed that after childbirth and at a distance from childbirth (V2) DUSP1 finds a rate like the control group (*p<0.0001*). Same modulation profile of FLT1 concentration was observed in PE donors compared to healthy donors (*p<0.0001*, all).

**Figure 5.**
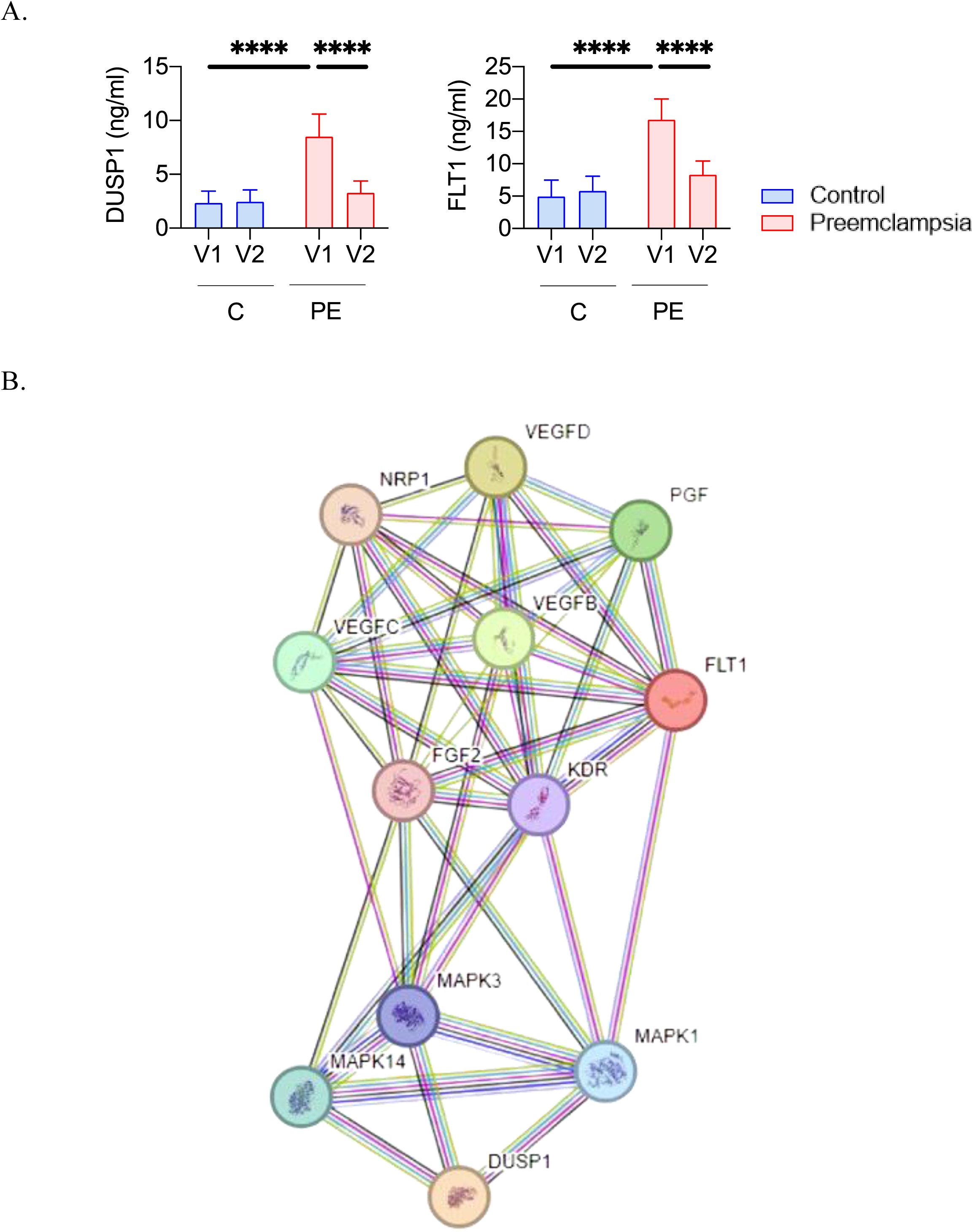
DUSP1 represents a biomarker candidate of preeclampsia. (**A**) Quantification of DUSP1 and FLT1 protein level by ELISA on the plasma (from V1 and V2) from 9 controls (C) and 9 preeclamptic women’s (PE). Statistical analysis was performed with one-way ANOVA and Tukey’s multiple comparison test. \**p*≤*0.05*, \*\**p*≤*0.01*, \*\*\**p*≤*0.001* and \*\*\*\**p*≤ *0.0001*. (**B**) Protein pathways linked to DUSP1.

Finally, to further understand molecular signature changes, including *DUSP1* gene, and how it might reflect PE pathophysiology, we performed a protein pathway analysis (**Figure 5B**). There were 12 proteins associated with DUSP1. Among them, we found FLT1 suggesting common pathways that can explain the same expression profiles. There were also proteins associated with VEGF (vascular endothelial growth factor) and PGF (platelet growth factor) previously described as associated with PE pathophysiology ^14^. All together these results highlighted a new blood biomarker candidate for diagnosis of PE in pregnant women.

## Discussion

The clinical diagnosis of PE remains challenging and is often delayed because of the lack of reliable early biomarkers. Although studies have used large biobanks and cohorts, the identification of efficient biomarkers for the early diagnosis of PE is still warranted. In this study, we adopted a specific study design strategy to investigate new candidate as biomarkers of PE by the evaluation of the expression of genes at the blood and placenta levels in women with PE, which were not found postpartum at distance of delivery. Our study highlights DUSP1 as a non-invasive blood biomarker candidate of PE.

Current screening tools are essentially in the form of a diagnostic tree combining several risk factors for PE to predict its occurrence in the short term. They combine several early markers: clinical (mean blood pressure), ultrasound (pulsatility index of uterine arteries) and biological (PAPP-A and PlGF), allowing to predict the risk of PE before term with a risk of false positive around 10 to 15% ^19,20^. Recent data from the literature open the prospect of a promising molecular approach in PE by studying gene expression in this pathology ^21–23^. However, studies focus on the investigation of genes on samples, at the blood or placenta level, only at the time of diagnosis. The strength of our study was primarily its prospective character, which made our study robust. Controls were rigorously matched to patients with PE on the 2 major confounding factors: maternal age and gestational age at diagnosis. The two groups (PE and control) were comparable across all initial characteristics, thus addressing potential confusion bias. The transversality of our study is also a strong point as patients in each group were followed from the first clinical manifestations of PE to postnatal. Thus, each patient was taken at the 3 major events of the pathology, which are: diagnosis (first symptoms), childbirth (signs of severity indicating fetal birth and/or maternal rescue), and postpartum (remission). This transversality is a major asset to follow the evolution of the transcriptional signature of the PE according to the evolution of the disease.

Our study highlighted a specific gene signature of PE. Among the modulated genes, the associated biological processes were previously described in the pathophysiology of PE ^24,25^. Interestingly, we were able to highlight the *FLT1* gene whose biomarker role was well-documented in PE ^13,26^. The presence of this gene indicates that the cohort choice and design strategy of the study is like previous studies. We show that *FLT1* has the same significant expression modulation profile as *DUSP1*, and both find a physiological basal state at distance from pregnancy. We also show that *FLT1* is present in the *DUSP1* pathway. Additional studies were carried out to define the relevance of DUSP1 and FLT1 in PE as an isolated biomarker candidate or as a signature.

Our study identified DUSP1 as a biomarker candidate of PE. DUSP1 is part of a large superfamily of 30 types of DUSP involved in signal transduction pathways that inactivated mitogen-activated protein kinases (MAP kinases). Specifically, DUSP1 modulation induces changes in several pathways such as MAP kinase phosphatase activity, tyrosine kinase receptor activity, angiogenesis and cell-cell-signaling ^27^. Its role as actors of tumor biology is well documented ^28^. Interestingly, several studies highlighted the relationship between DUSP1 and hypoxia, a major contributor to the abnormalities reported in the placenta from PE women. Hypoxic conditions lead to *DUSP1* over-expression associated to an increased interaction with the hypoxia-inducible factor (HIF)-1 alpha unit ^29^ involved in the pathogenesis of PE ^30^. Further studies are needed to highlight the mechanism of action of DUSP1 in PE.

Previous study has investigated DUSP1 for the identification of PE ^31^. The authors investigated *DUSP1* expression from the placenta tissue avec from blood cord. The authors report confusing data concerning the expression of *DUSP1* in placental tissue. *DUSP1* mRNA expression from the PE group was found to be significantly lower than that of the healthy group. In contrast, protein levels, evaluated by immunohistochemistry, were similar in the PE and control groups. Considering DUSP1 as biomarker the authors investigated DUSP1 protein levels from blood cord. They reported significant lower DUSP1 expression in PE women compared to healthy donors. Moreover, the authors used a limited cohort (400 controls *versus* 5 PE samples) and did not investigate the gestational age as diagnostic that constitutes a major confounding factor associated with potential confusion bias. In contrast, Yonghong Wang et al. have reported an indirect role for DUSP1 in the occurrence of the PE ^32^. The authors reported that miR-141-5p reduced *in vitro* DUSP1 expression affecting MAPK/ERK pathway to promote the PE. Although further studies are needed to identify the role of DUSP1 in PE, this study revealed the expression of DUSP1 by immortalized JEG-3 trophoblastic cells whose role in pregnancy and its involvement in the occurrence of PE remain to be defined.

In conclusion, based on an original study design, our study reports genes associated with PE which some of them were previously described associated with the pathophysiology of PE. The investigation of DUSP1 on a larger cohort before and after the first events of PE, including the severity aspect of the disease, is necessary to confirm its value as a biomarker. RANSPre study, a French multicenter study, could be an alternative strategy to evaluate this candidate.

## Author contribution

J.A and A.D; and S.M, J.L.M and F.B contributed equally to this work. J.A and A.D were involved in experimental design and execution, data analysis and figure generation. A.D and J.F.C collected samples used in these experiments and participate to experiments. G.C and J.P realized RNAseq data analysis and figure generation. A.D and F.B initiated the clinical study and participated in the analysis of the clinical characteristics of the cohort. N.R carried out the statistical analyses. S.M, J.L.M and F.B contributed expertise in the guidance for experimental design, supervised the study and manuscript preparation and mentorship.

## Acknowledgments

This work was supported by the French government under the Investissements d’avenir (Investments for the Future) program managed by the Agence Nationale de la Recherche (reference number 10-IAHU-03). This work was supported by the “Comité” 10 28 project managed by the “Assistance Publique Hopitaux de Marseille” (reference ID RCB 2012-A00633-36).

---

**Table S1.**
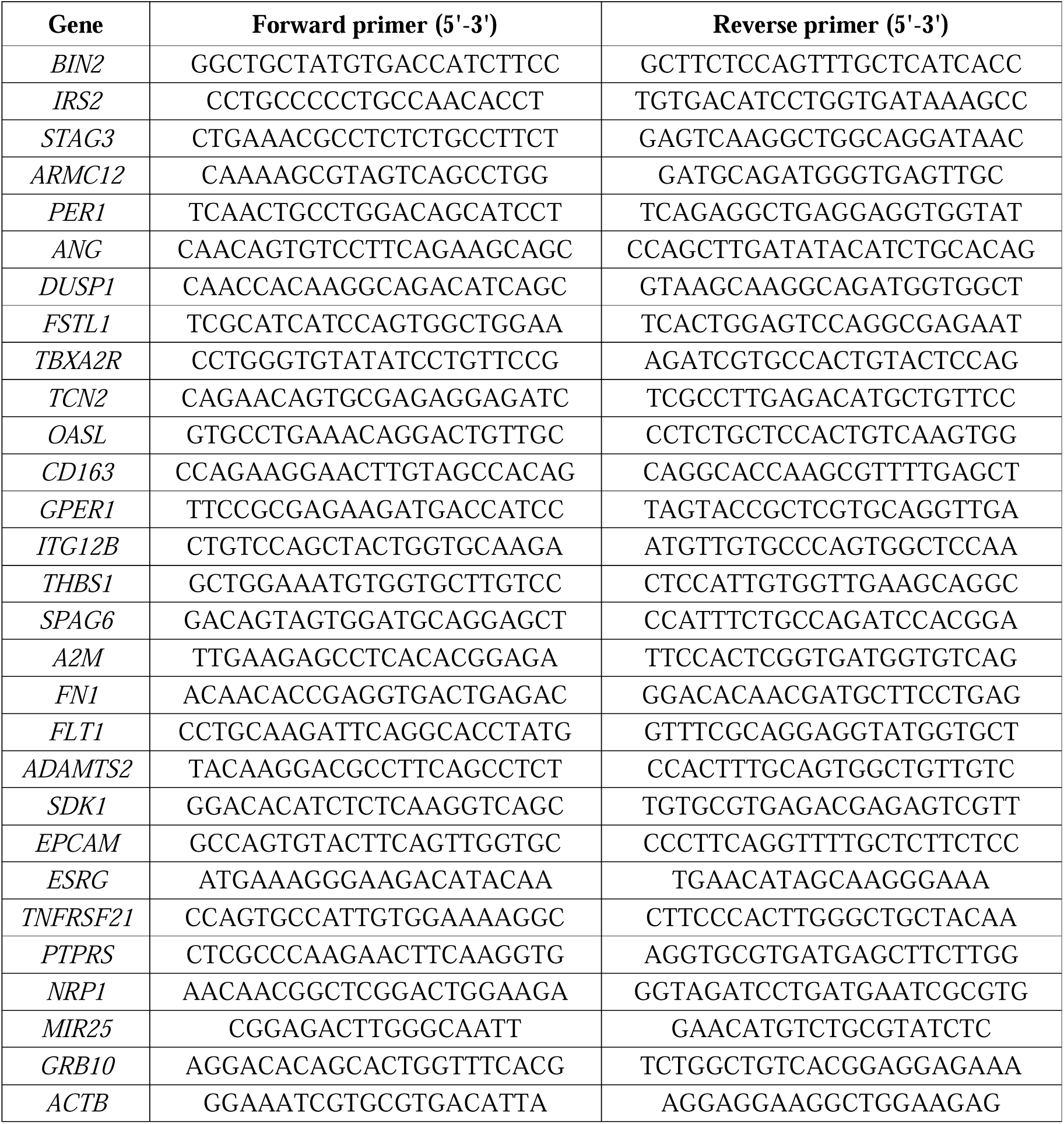
List of primers used for qRTPCR.

**Table S2.** List of genes associated to the preeclampsia signature.

| **Clusters** | **Gene name** | **Complete name** |
| --- | --- | --- |
| **Angiogenesis,**<br>**Differentiation** | *ARMC12* | Armadillo repeat containing 12 |
|  | *ANG* | Angiogenin |
|  | *FSTL1* | Follistatin like 1 |
|  | *DUSP1* | Dual specificity phosphatase |
|  | *FN1* | Fibronectin |
|  | *FLT1* | FMS related receptor tyrosine kinase 1 |
|  | *NRP1* | Neuropilin 1 |
|  | *GRB10* | Growth factor receptor bound protein 10 |
| **Cell**<br>**adhesion** | *ITGA2B* | Integrin subunit alpha 2b |
|  | *THBS1* | Thrombospondin |
|  | *SDK1* | Sidekick cell adhesion molecule 1 |
|  | *EPCAM* | Epithelial cell adhesion molecule |
|  | *PTRS* | Protein tyrosine phosphatase receptor type S |
| **Cell cycle** | *IRS2* | Insulin receptor substrate 2 |
|  | *STAG3* | Stromal antigen 3 |
|  | *MIR25* | Micro RNA 25 |
|  | *ESRG* | Embryos stem cell related |
|  | *GPER1* | G protein coupled estrogen receptor 1 |
|  | *PER1* | Period circadian regulator 1 |
|  | *BIN2* | Bridging integrator 2 |
| **Inflammatory**<br>**response** | *CD163* | Cluster differentiation 163 |
|  | *TNFRSF21* | TNF receptor superfamily member 21 |
| **Cell**<br>**metabolism** | *TCN2* | Transcobalamin 2 |
|  | *SPAG6* | Sperm associated antigen 6 |
|  | *A2M* | Alpha 2 macroglobulin |
|  | *ADAMTS2* | ADAM metalloproteinase with thrombospondin type 1 motif 2 |

## Supplemental figures legends

**Figure S1.**
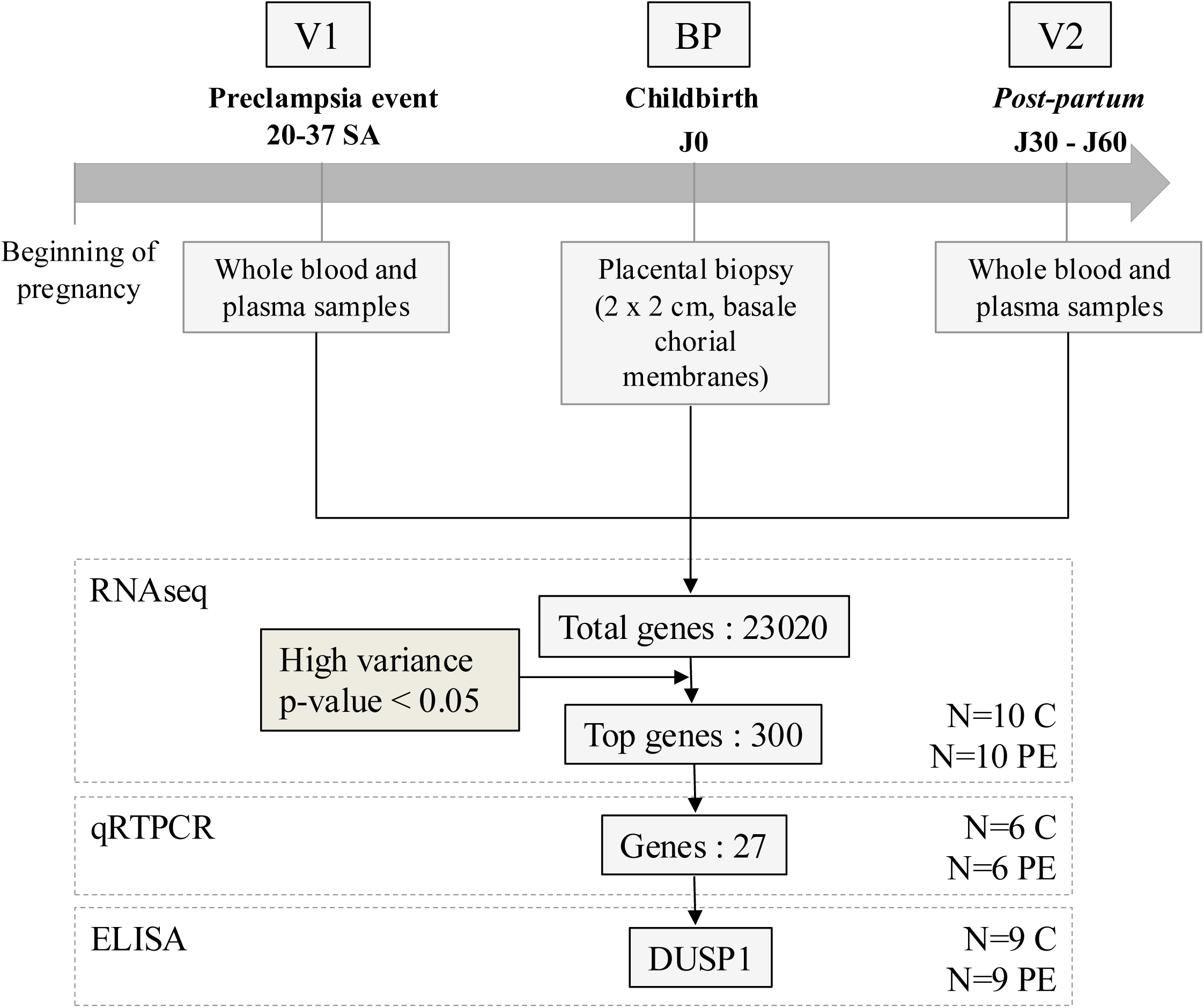
Study design. Samples from included patients were taken at the 3 major events: at the diagnosis of preeclampsia (PE) (V1, first symptoms), childbirth (Biopsy from placenta (BP), signs of severity indicating fetal birth and/or maternal rescue), and postpartum (V2, remission). V1 and V2 corresponded to whole blood and plasma samples and BP for biopsy of placenta. Total ARN were isolated and analyzed through an RNAseq method (10 samples for PE and control groups). qRTPCR (6 samples from PE and control (C) groups) and ELISA (9 samples from PE and control groups) were realized on isolated RNA and plasma samples, respectively. RNAseq, RNA sequencing; qRT-PCR, Quantitative reverse transcription-polymerase chain reaction; ELISA, enzyme-linked immunosorbent assay.

## References

1. Dimitriadis, E. et al. Pre-eclampsia. Nat. Rev. Dis. Primer 9, 1–22 (2023).

2. Chappell, L. C., Cluver, C. A., Kingdom, J. & Tong, S. Pre-eclampsia. The Lancet 398, 341–354 (2021).

3. Ives, C. W., Sinkey, R., Rajapreyar, I., Tita, A. T. N. & Oparil, S. Preeclampsia— Pathophysiology and Clinical Presentations: JACC State-of-the-Art Review. J. Am. Coll. Cardiol. 76, 1690–1702 (2020).

4. Tita, A. T. et al. Treatment for Mild Chronic Hypertension during Pregnancy. N. Engl. J. Med. 386, 1781–1792 (2022).

5. Chappell, L. C. et al. Planned early delivery or expectant management for late preterm pre-eclampsia (PHOENIX): a randomised controlled trial. Lancet Lond. Engl. 394, 1181– 1190 (2019).

6. Say, L. et al. Global causes of maternal death: a WHO systematic analysis. Lancet Glob. Health 2, e323–e333 (2014).

7. Dimitriadis, E. et al. Pre-eclampsia. Nat. Rev. Dis. Primer 9, 8 (2023).

8. Redman, C. W. G., Staff, A. C. & Roberts, J. M. Syncytiotrophoblast stress in preeclampsia: the convergence point for multiple pathways. Am. J. Obstet. Gynecol. 226, S907–S927 (2022).

9. Zhang, R. et al. PD-L1 enhances migration and invasion of trophoblasts by upregulating ARHGDIB via transcription factor PU.1. Cell Death Discov. 8, 1–10 (2022).

10. Yin, W. et al. Genetic Variation in ANGPTL4 Provides Insights into Protein Processing and Function *. J. Biol. Chem. 284, 13213–13222 (2009).

11. Meinhardt, G. et al. Pivotal role of the transcriptional co-activator YAP in trophoblast stemness of the developing human placenta. Proc. Natl. Acad. Sci. 117, 13562–13570 (2020).

12. MacDonald, T. M., Walker, S. P., Hannan, N. J., Tong, S. & Kaitu’u-Lino, T. J. Clinical tools and biomarkers to predict preeclampsia. EBioMedicine 75, 103780 (2021).

13. Verlohren, S. et al. Clinical interpretation and implementation of the sFlt-1/PlGF ratio in the prediction, diagnosis and management of preeclampsia. Pregnancy Hypertens. 27, 42– 50 (2022).

14. Zeisler, H. et al. Predictive Value of the sFlt-1:PlGF Ratio in Women with Suspected Preeclampsia. N. Engl. J. Med. 374, 13–22 (2016).

15. Rolnik, D. L. et al. ASPRE trial: performance of screening for preterm pre[eclampsia. Ultrasound Obstet. Gynecol. 50, 492–495 (2017).

16. Rolnik, D. L. et al. Aspirin versus Placebo in Pregnancies at High Risk for Preterm Preeclampsia. N. Engl. J. Med. 377, 613–622 (2017).

17. Love, M. I., Huber, W. & Anders, S. Moderated estimation of fold change and dispersion for RNA-seq data with DESeq2. Genome Biol. 15, 550 (2014).

18. Mezouar, S. et al. Full-term human placental macrophages eliminate *Coxiella burnetii* through an IFN-γ autocrine loop. Front. Microbiol. 10, 2434 (2019).

19. O’Gorman, N. et al. Competing risks model in screening for preeclampsia by maternal factors and biomarkers at 11-13 weeks gestation. Am. J. Obstet. Gynecol. 214, 103.e1–103.e12 (2016).

20. Chappell, L. C. et al. Diagnostic accuracy of placental growth factor in women with suspected preeclampsia: a prospective multicenter study. Circulation 128, 2121–2131 (2013).

21. Enquobahrie, D. A. et al. Differential placental gene expression in preeclampsia. Am. J. Obstet. Gynecol. 199, 566.e1–566.11 (2008).

22. Okazaki, S. et al. Placenta-derived, cellular messenger RNA expression in the maternal blood of preeclamptic women. Obstet. Gynecol. 110, 1130–1136 (2007).

23. Textoris, J. et al. Evaluation of Current and New Biomarkers in Severe Preeclampsia: A Microarray Approach Reveals the VSIG4 Gene as a Potential Blood Biomarker. PLoS ONE 8, e82638 (2013).

24. Kondoh, K., Akahori, H., Muto, Y. & Terada, T. Identification of Key Genes and Pathways Associated with Preeclampsia by a WGCNA and an Evolutionary Approach. Genes 13, 2134 (2022).

25. Mohamad, M. A. et al. A Review of Candidate Genes and Pathways in Preeclampsia-An Integrated Bioinformatical Analysis. Biology 9, 62 (2020).

26. Stepan, H., Hund, M. & Andraczek, T. Combining Biomarkers to Predict Pregnancy Complications and Redefine Preeclampsia: The Angiogenic-Placental Syndrome. Hypertens. Dallas Tex 1979 75, 918–926 (2020).

27. Huang, C.-Y. & Tan, T.-H. DUSPs, to MAP kinases and beyond. Cell Biosci. 2, 24 (2012).

28. Shen, J. et al. DUSP1 inhibits cell proliferation, metastasis and invasion and angiogenesis in gallbladder cancer. Oncotarget 8, 12133–12144 (2017).

29. Liu, C. et al. Dual-specificity phosphatase DUSP1 protects overactivation of hypoxia-inducible factor 1 through inactivating ERK MAPK. Exp. Cell Res. 309, 410–418 (2005).

30. Tal, R. The role of hypoxia and hypoxia-inducible factor-1alpha in preeclampsia pathogenesis. Biol. Reprod. 87, 134 (2012).

31. Wang, J., Zhou, J.-Y., Kho, D., Reiners, J. J. & Wu, G. S. Role for DUSP1 (dual-specificity protein phosphatase 1) in the regulation of autophagy. Autophagy 12, 1791–1803 (2016).

32. Wang, Y. et al. miR-141-5p regulate ATF2 via effecting MAPK1/ERK2 signaling to promote preeclampsia. Biomed. Pharmacother. 115, 108953 (2019).

